# Lack of early animal fossils: insights from taphonomic experiments on placozoans and their traces

**DOI:** 10.1101/2023.08.02.551688

**Authors:** E.B. Naimark, Yu.V. Lyupina, M.A. Nikitin, A.D. Finoshin

**Affiliations:** Borissiak Paleontological Institute, Russian Academy of Sciences, 123 Profsouznaya St, Moscow, Russia; Koltsov Institute of Developmental Biology, Russian Academy of Sciences, 26 Vavilov St, Moscow, Russia; Belozersky Institute of Physicochemical Biology, Lomonosov Moscow State University, Leninskye gory 1 building 40, Moscow, Russia

## Abstract

Placozoa are simple two-layered multicellular animals. Can such a simple animal preserve in the fossil record? The results of our taphonomic experiments showed that it is chemically possible; however, the chances of the presevation are negligible for two reasons. Firstly, the resistance of living *Trichoplax* to lethal factors turned out to be quite high. Secondly, post-mortem changes in the body of *Trichoplax* preclude preservation. We observed these changes for the first time: their bodies immediately disintegrated into a cloud of individual dead cells, apparently unrecognizable in the fossil record. We hypothesized that the absence of a basement membrane, which unites each cell layer into the epithelia, is a key factor in this dramatic disintegration. This hypothesis adds an option to the explainations of the gap between the time of emergence of metazoan body fossils and the reconstructed time of metazoan divergence from a single-celled ancestor. However, it is theoretically possible to find ichnofossils of *Trichoplax*-like locomotion. For the first time, we observed traces of *Trichoplax* on a soft substrate and identified a previously unknown type of traces. They were 3D chains of compacted round lumps of the substrate, similar in some features to the Precambrian meniscus ichnofossils.

## INTRODUCTION

The fossil record of Metazoa are known from the late Precambrian (around 600 Ma), while molecular age of their origin was estimated to be rather old to be around 800-900 Ma [1]. Currently, the main explanation for this gap is that the ancestors of metazoans were too small, thin, and soft to become a fossil and to be found by paleontologists [2]. Contrary to this explanation – tiny/ thin/ soft – unicellular fossils do exist, and also fossils of soft medusa-like animals exist indicating that tiny and thin animals could form a fossil and such fossils could be found in deposits. Thus we possibly omit something important in the discussion on the earliest Metazoans fossils formation. Taphonomic experiments help to understand what a particular type of animal would look like in a fossil, but nothing is currently known about the preservational potential of early metazoans. Due to the lack of such knowledge, we cannot discuss whether the gap between the molecular age of the origin of the Metazoa and their fossil record is taphonomic in nature.

Here we focused on the Placozoans with their very simple morphology to explore the possibility of preservation in the earliest Metazoans. There are no experimental data on preservational potential of placozoans. Also, to our surprise, there was not even any information about how *Trichoplax* dies and how its postmortem transformation proceeds to be snapshotted as a fossil. In this work, we filled this gap.

All known Placozoa [3–5] are multicellular animals with an extremely simple body plan. Their body consists of ventral and dorsal cell layers, joined at the edges by a narrow band of generative cells [6–10]. The connection of cells – *i*.*e*. the basis of multicellularity – is mediated by simple adhaerens junctions, while they do not have any tight or septate junctions [11]. The extracellular matrix consists mainly of mucin, while collagen IV and laminin, the main building blocks of basement membranes, are absent in the cell layers. Moreover, there are also no a basal membrane underlain the cell layer [9].

An animal with such a simplified, albeit recurrent, organization of multicellularity can serve as a taphonomic model for early metazoans that did not yet have fully evolved cell junctions, an extracellular matrix and basement membranes [12–16].

Our study answers the question of what might be left of a *Trichoplax*-like animal in the fossil record. Both body remains and traces can be preserved: the former as body fossils and the latter as ichnofossils. Accordingly, our work is divided into two parts: the first part is devoted to the possible preservation of body remains, and the second part relates to Placozoan traces. It should be noted that, along with the absence of data on the postmortem transformations of *Trichoplax* body, neither does data on its locomotion on soft substrate exist, though it is on soft sediment that an animal can leave a trace noticeable to a paleontologist. Therefore, in both cases, this work pioneered the taphonomic studies of Placozoans.

The objectives of this study were as follows:

- to study the deposition of preservative cations on the *T. adhaerens* dead bodies which necessarily preceding fossilization. This task relates to the capacity of adhesive molecules in multicellular tissues to effectively bind to preservative cations (Al-ions) [17], this process facilitates the fast preservation and fossilization of decaying bodies. However, the adhesive complex in Placozoa, as it was mentioned above, is depleted compared to other multicellular animals. Thus the doubts exist if the adhesive complex of placozoans is enough to allow for preservation of their body remains.
- to study the body transformation of *Trichoplax* after death caused by various lethal factors: UV (physical factor), the high concentrations of sodium azide (chemical factor); and next, to reveal the similarity between these postmortem transformations in order to elicit the possible features of placozoan-like fossils.
- to study the survival of *Trichoplax* under the conditions accompanying the catastrophic burial of soft-bodied animals, namely, deposition under a layer of fine-grained sediment, acidification in organic-rich fine-grained sediment, hypoxia and hydrogen sulfide contamination (for review see [18,19]). Are these conditions actually lethal for *Trichoplax*?
- to study the locomotion of *Trichoplax* on the surface of fine-grained sediment.

## MATERIALS

In our experiments, we used the culture *T. adherens* from the Belozersky Institute of molecular biology (this culture originates from B. Schierwater placozoan collection). The culture was maintained in artificial sea water (ASW) on dishes with *Tetraselmis marina* (Chlorophyta) or *Oscillatoria* (Cyanobacteria) as a food source. When growing on *Oscillatoria*, the *Trichoplax’* bodies become pinkish as they do not digest red pigment, resulting in its accumulation in cells of the body. This property proved useful in experiments on locomotion because it helped to visualize the *Trichoplax’* bodies against a white substrate. It should be noted that common vital dyes (tolluidine, Bengal rose, acridine orange) appeared to be lethal for *Trichoplax* and kill them in first hour (though vital staining has been used for *Trichoplax* by other authors [3]).

Kaoline with the kaolinite content around 97% was used as a fine-grained substrate with particles less than 1 μm [18]. Kaolinite has been shown to slow down organic decay and therefore it may contribute to preservation of soft-bodied organisms in sediment [19–21].

## METHODS

Here we describe the methods in an order corresponding to the logical series of the experiments and the subsequent results.

The deposition of preservative cations (namely Al^3+^) was detected utilizing the specific fluorescent dye lumogallion [19]. Lumogallion (5-chloro-3[(2,4-287 dihydroxyphenyl)azo]-2-hydroxybenzenesulfonic acid) responds to bound Al in low quantities by emitting a green light under UV excitation (see [22] for technical details). The *Trichoplax* bodies that were killed by adding methanol into the small Petri dish were incubated for 30 minutes in the 0,001M alum solution (KAl(SO4)2, and fixated with 4% PAF and then rinsed from the alum remains by distilled water to leave only Al bound to organic tissues. The samples prepared in this way were incubated with lumogallion for 1 hour at 50°C, rinsed and set up for microscopy (under confocal fluorescent microscope Carl zeiss LSM 880 airyscan and fluorescent microscope Leica DM RXA2 in Institute of developmental biology). We kept samples for no more than 8 hours at 4°C. The Al-negative control (a sample without incubation in the alum solution) and Al-lumagallion negative control (autofluorescence) were compared with the experimental samples.

The postmortem transformations were recorded in a micro-camera of a depth of around 0,5 mm filled with a solution of sodium azide (0,4 and 0,1%) in the first experiment and with ASW in the second experiment. In the former one, we also added eosin into the solution in order to test cells on a “live or dead” condition. Eosin does not dye live cells, but dead cells become pink as the cellular membrane is destroyed and the dye binds to cytoplasmic proteins. This test allowed us to distinguish the death of single cells from the death of an entire multicellular organism. The eosin itself appeared to be lethal, but in the low concentration we applied, its lethal action manifested in 20-24 hours as we could observe in the supplemental tests. In the second experiment, the *Trichoplax* in the micro-camera was radiated by UV, which is known to be detrimental to it. For visualization of the cells, we stained live *Trichoplax* with nuclear fluorophore DAPI, which emitted in blue range when bound with DNA.

The uniformly designed experiments were set to study the survival of *Trichoplax* in lethal burial conditions. Several *Trichoplax* specimens (5-16) were resettled to small Petri dishes with small amounts of ASW; after they attached to the bottom we gently added the solutions according the experimental goal of each experiment. The experimental dishes did not contain algae thus *Trichoplax* did not feed throughout the experiments.

When testing the survival of *Trichoplax* in decreased acidity, the pH of the experimental ASW was lowered from the initial value of 7.8 to 5.0, 6.0 and 7.0 by adding HCl. Such a pH range characterizes the acidity in fine-grained sediments with decaying organics [19,23,24].

Tests on survival in hydrogen sulfide contamination were set up in ASW, containing 50, 100, and 300 mg/L of S^2-^. The latter concentration of 300 mg/L characterizes the level of hydrogen sulfide in the Black Sea bottom sediments; the vital activity of multicellular animals there is strongly, although not completely suppressed [25].

The solutions were prepared from the corresponding weighted portions of Na_2_S * 9H_2_O. The survival of *Trichoplax* in high H_2_S we tested in 50-ml horizontal flasks (Fig. 6, inset) tightly sealed to prevent any escaping of the hydrogen sulfide.

It seems that the experiments on hydrogen sulfide contamination should correlate with the experiments on hypoxia. However, H_2_S itself can be fatal instead of deficiency of oxygen. We have provided additional tests for hypoxia by directly restricting oxygen respiration. In these tests, we used sodium azide. In all organisms, this substance inhibits cytochrome oxidase and catalase, which mediate cellular oxygen respiration. We used sodium azide concentrations of 0.005, 0.01, 0.02, 0.1, 0.2, and 0.4%. We chose this range considering that the lethal range for bacteria is 0.01-0.02% and about 0.05% for unicellular amoebae.

In all experiments, we observed and imaged the specimens under a light microscope, recording the moments when each specimen exhibited locomotion, stopped, and disintegrated. In some cases, we video-recorded single specimens.

In the experiment imitating catastrophic deposition, 10 animals (pigmented or white) attached to the Petri dish’ bottom were poured over by a thick suspension of kaoline, with the depth of the sediment after settlement being about 2.5-3 mm. We made two repetitions where in the first one we videotaped the *Trichoplax* by inverted microscope from the bottom side of the dish, while in the second dish we registered the animals on the upper side of the sediment. Thus, the first repetition shows the behavior of *Trichoplax* under the layer of the sediment, and the second repetition shows the number of animals which appeared on the sediment surface (which means that they survived after burying). Also, we could register the traces of the animals after they got to the surface.

When studying traces, we placed pigmented animals on the kaoline sediment after the kaoline suspension had been settling for three hours or had been centrifugating on low speed for consolidation. We videotaped the locomotion of single specimens for 2.5-3 hours, and after 20-24 hours the resulting traces were imaged. The traces appeared to be very unstable and thus they were easily destroyed in any agitation. Therefore, in order to retain the shape of the traces, the dishes with animals stayed unmoved on the microscope table throughout the entire experiment.

For the experiment on feeding on bacterial mats, we grew *Oscillatoria* on the thinly grounded limestone for two-three days. In this way, we got the sediment glued by the bacterial film.

## RESULTS

### Deposition of Al-ions on dead bodies of *Trichoplax*

The dead bodies of *Trichoplax* effectively adsorbed Al-ions which were clearly recognized by the bright green fluorescence specific for bound Al. This sharply contrasts with the very weak fluorescence in the Al-negative control and autofluorescence (Fig. 1). We concluded from this result that dead bodies of animals with the same rudimentary set of adhesive complex and extracellular matrix as in *Trichoplax* can, in principle, be naturally preserved in aluminosilicate sediment.

**Fig.1.**
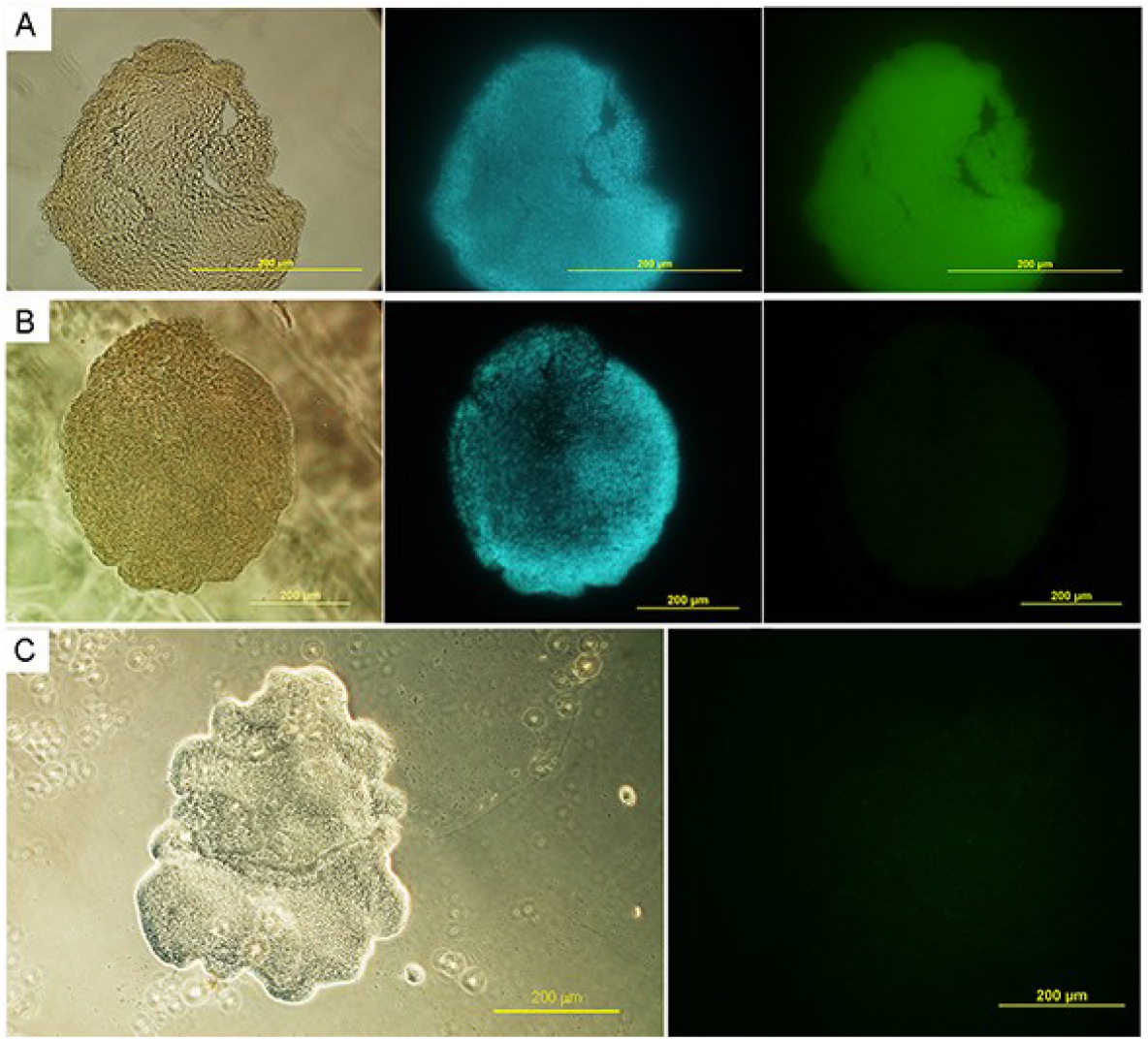
Staining of dead *Trichoplax* bodies with lumogallion. A. the experimental sample incubated in the alum solution (see Methods): left – under a light microscope, middle – under UV-excitation showing blue DAPI-stained cell nuclei, right – under excitation at 488 nm and emitting at 560-570 nm (green) specific for the bound Al-ions. When imaging the fluorescence of the samples, the exposure time was the same for the experiment and controls to allow the comparison of the intensity of the emitting. B. the same for the control sample which was incubated without the alum solution. C. the same for autofluorescence; (DAPI staining is omitted here).

### Post-mortem transformations of a *Trichoplax* body

This paragraph describes the post-mortem transformations of a *Trichoplax* body, answering the question what kind of placozoan remains a paleontologist can expect to find in the fossil record. We studied these transformations in two unequivocally lethal factors, which are completely different in their action: hypoxia caused by a very high concentration of sodium azide (0.4%) and UV radiation.

In sodium azide, the order of transformation of *Trichoplax* bodies was as follows (Fig. 2). After the exposition in azide, the locomotion of the live Trichoplax stopped from between 1 to 2 hours varying among individual; then their bodies contracted slightly, after which they relaxed and started increasing again. Simultaneously or slightly after this, a small spot of dead cells formed. The death of single cells could be evidenced by their change in color from whitish to pink in the presence of eosin in the experimental solution. Very quickly after the pink spot appeared, individual dead (pink) cells began to separate from the dead spot one after another; this movement resembled thin streams (arrow in Fig. 2).

**Fig.2.**
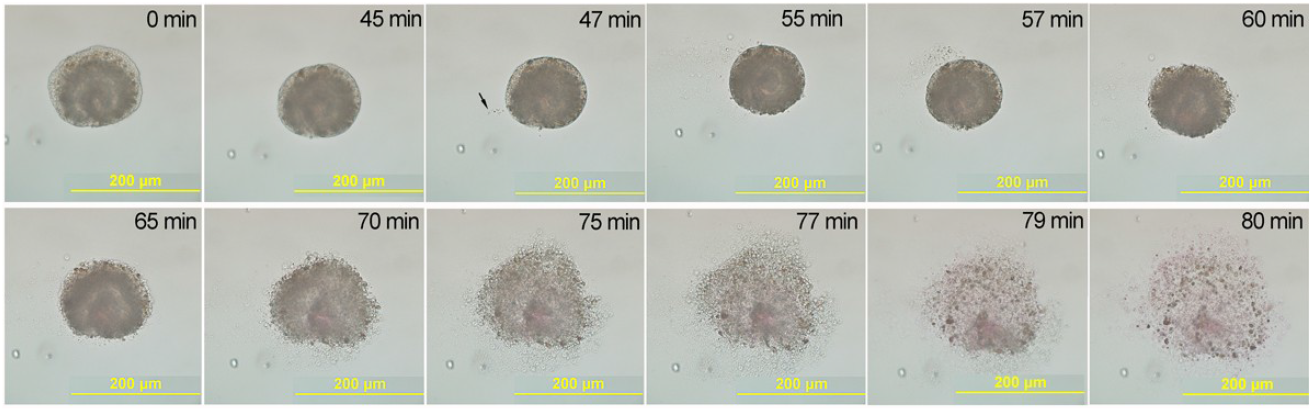
Postmortem transformation of the *Trichoplax’* body under sodium azide, concentration 0.4%; eosin staining; the times indicate the beginning of observation which started one hour after the experiment began. In the first hour (not shown here), the animal moved, then it stopped and began to shrink; the formation of a pink spot of dead cells in the center of the body can be seen. Then, on a certain place on the edge of the body, the cells began to detach from the edge and move to the side (arrow); after five minutes the process of cell detachment covers the entire edge of the body, and over the next 20 minutes, the dead cells scattered as a cloud. See video 1.

In a short time, the cells began to detach and floated away over the entire perimeter of the body. By this time, all the detaching cells had died, and the media around the body had acquired a pinkish color. In the next five to ten minutes, the entire body was collapsing; after fifteen minutes, all the cells in the body had died, and formed a cloud of individual dead cells and cellular remains. It is worth noting that the experimental dishes stayed immobile, so neither oscillation nor micro-currents in the dishes could move the dead cells. The dead cells were dispersed due to the physicochemical interactions of their surface molecules with salt water.

The same pattern of lethal transformations was also observed under the action of UV (Fig. 3). Three minutes after the start of irradiation, the animals stopped moving, including the beating of cilia. Two to three minutes later, they contracted slightly, reducing their surface area by 15-20%. After that, within one or two minutes, the body again relaxed, its area increased, and from one edge individual cells began to detach one after another, quickly moving away from the edge. The process of detachment of cells propagated over the entire body in two to three minutes, and the cells spread to the sides in a cloud.

**Fig.3.**
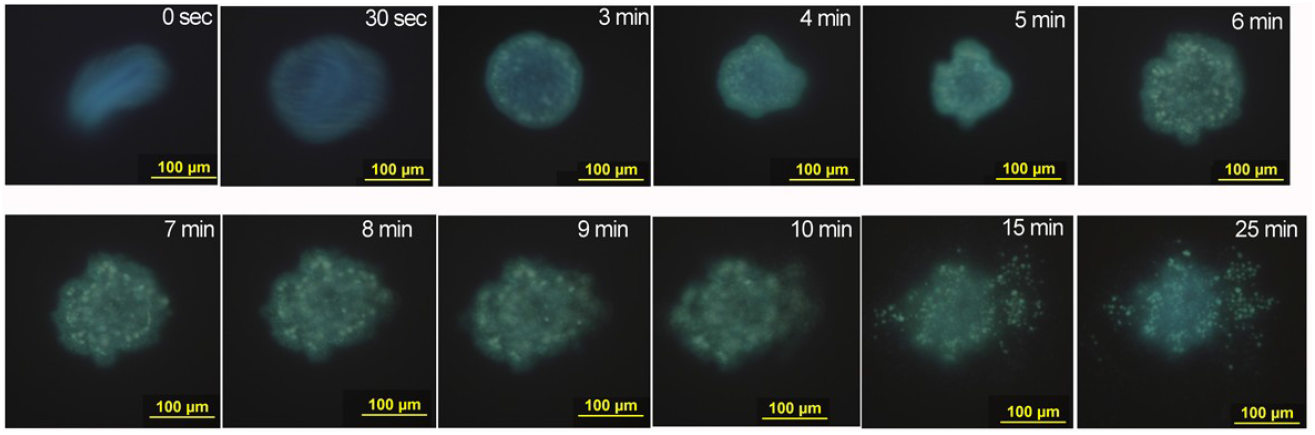
Post-mortem transformation of the body of *Trichoplax* under UV illumination; DAPI stain. In the first minute or two, the animal moved (because of the fast movement the two first photos are fuzzy), then it stopped, and began to shrink. After 5-6 minutes the body relaxed, increasing the surface area; then in one place the cells began to detach from the edge and move to the side, very quickly the process of cell detachment propagated over the entire body, and within 20 minutes the cells were dispersed in the shape/form of a cloud. See video2.

### Stability of *Trichoplax* under the influence of lethal burial factors

As shown by previous experiments, the criteria of *Trichoplax* death are the cessation of locomotion (early sign), body compression (intermediate sign) and disintegration into cells (late sign). Focusing on the early and late criteria, we tracked the stability of *Trichoplax* under the onset of unfavorable conditions that usually operate during a catastrophic burial: oxygen deficiency, hydrogen sulfide contamination, burying under a layer of fine-grained sediment, and the acidification of the environment.

#### *Reactions of Trichoplax to oxygen deficiency* (Fig. 4)

At high level of oxygen deficiency (i.e. high concentration of azide), the animals stopped moving during the first hour, during the next hour and a half they disintegrated into cells (Fig. 2).

**Fig.4.**
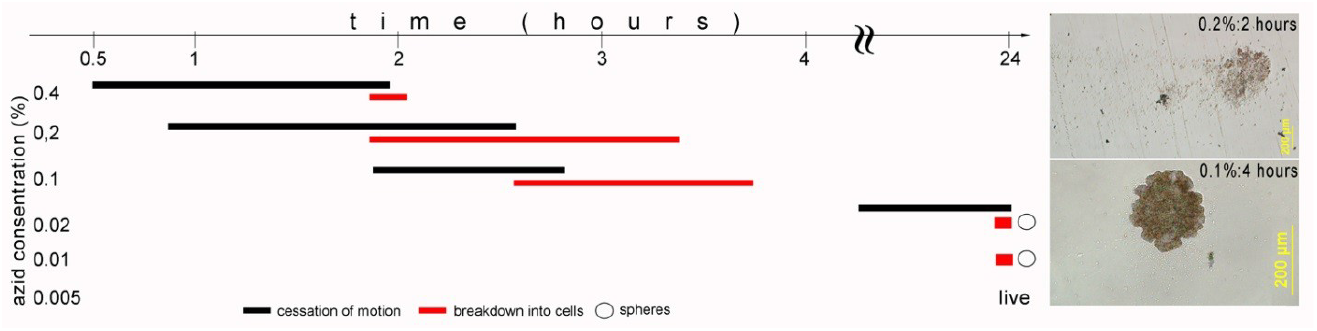
Trichoplax reactions in a series of concentrations of sodium azide. The bars show the first and last reactions among the group of individuals in each experimental dish (6-15 individuals): black bars – cessation of motion, red bars – disintegration of cells. For concentrations of 0.02-0.005, no changes were observed during the first 6 hours; changes registered after 24 hours. The insets on the right show the bodies’ transformations at two concentrations of azide, giving an expression of the postmortem patterns similar to those in Fig. 2, but differing in the speed of the process.

At a concentration of azide at 0.1% (ten times higher than bactericidal), the movement of *Trichoplax* was maintained for two hours, the deterioration of the integrity of the body and disintegration into cells began 2.5 hours after the start of the azid exposure. At a concentration close to the bactericidal one (0.01-0.02%), only one animal stopped moving and only after 4.5 hours of observation. All the rest did not change their motor activity during 6 hours of observation. However, after 24 hours at these concentrations, they still died, disintegrating into cells. At the same time, single living mobile spheres remained in both experimental dishes, which is a specific form of an asexual reproduction of *Trichoplax* [26] (Fig. 5A). At high concentrations of azide (0.1% and above), these spheres very quickly disintegrated into individual cells (Video 3).

**Fig. 5.**
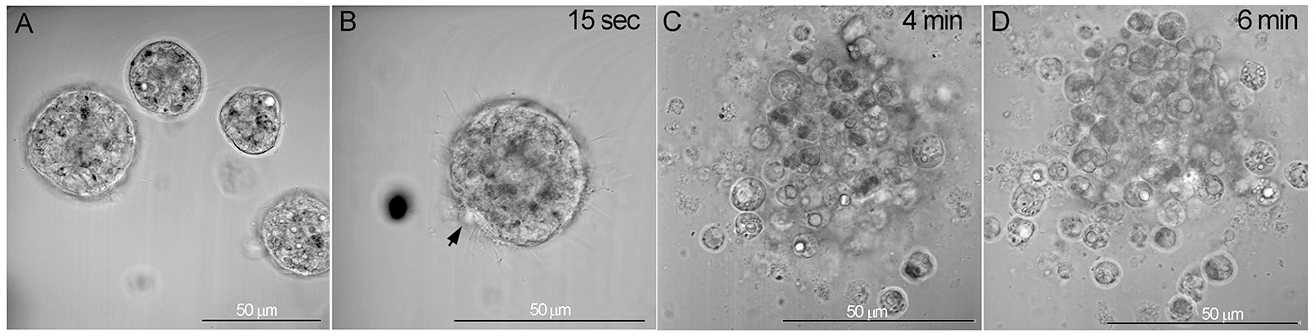
The numerous spheres appeared in unfavorable conditions. A. the spheres in the experimental dish at pH 6. B–D. disintegration of the spheres after adding the azide in a high concentration (0.4%): the tegument destroyed in one place (arrow) in 15 seconds in azide (B); in next 2-6 minutes the entire tegument destroyed and cells went out (C) and floated away (D).

At an azide concentration two times lower than the bactericidal one (0.005%), no signs of the inhibition of locomotion were observed, and after 24 hours all individuals in the dish remained mobile.

*At a hydrogen sulfide concentration of 300 mg/L*, all animals stopped moving after 20-25 minutes and disintegrated into cells within the first hour. At a concentration of 100 mg/l, the cessation of motor activity was recorded during the second hour, and throughout the third hour, all individuals disintegrated into cells. It is interesting to note that at this concentration, the asexual sphere had already begun to form on the dorsal side of one of the animals after only five minutes of exposure.

With a moderate hydrogen sulfide contamination of 50 mg/L, after 2 hours of observation, 1 animal out of 12 disintegrated into cells, after 3 hours all individuals stopped moving, and after the following hour, all individuals except for two disintegrated into cells. The remaining two individuals disintegrated over the next 6 hours. One movable sphere remained in the dish, however it disintegrated during the next day.

#### Decreased acidity

Given the range of acidity in an organic-rich marine sediments, we exposed the *Trichoplax* to an acidity pH 5.0, 6.0, and 7.0. In an acidity of pH 5, the locomotion of *Trichoplax* (9 specimens) did not change throughout the 8 hours of observation. At the same time, the asexual spheres began to form after two hours of exposition. In the next day (18-20 hours), one animal disintegrated into cells, and a significant number of the spheres appeared in the experimental dish (Fig. 5A). The spheres can actively move.

At pH 6, there was also no inhibition of locomotion; the same as at pH 5.0, we noted numerous live spheres at the next day. At pH 7, neither changes in locomotion nor any spheres were observed. Therefore, the formation of the spheres was induced by acidity below pH 6.

#### Burying under thick layer of sediment

When buried under a layer of sediment, *Trichoplax* behaved in two ways. In the first scenario, *Trichoplax* remained under the layer of sediment. It continued to actively move, forming passages between sediment particles. Its viability was also evidenced by the fact that it was capable of division: after 7.5 hours in the sediment, *Trichoplax* divided; both daughter individuals remained active until the end of observations (two days) (Fig.7 and Video 4).

**Fig.6.**
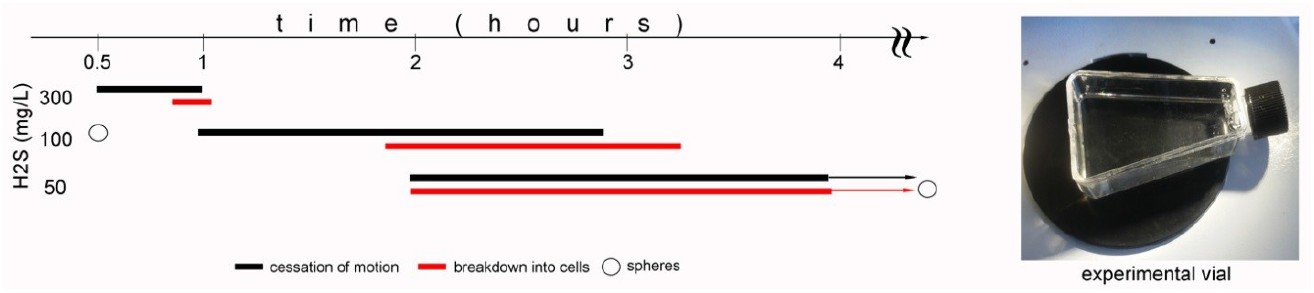
*Trichoplax* reactions under the action of hydrogen sulfide. The columns are the same as in Fig. 4 (9, 11, and 12 individuals, respectively, for concentrations of 50, 100, and 300 mg/l); for a concentration of 50 mg/L, observations of the last two live animals were stopped after 4 hours. They survived the next day (24 hours). The inset on the right shows a photograph of the vial used in this experiment; it does not distort the view, allowing to observe animals under a microscope.

**Fig.7.**
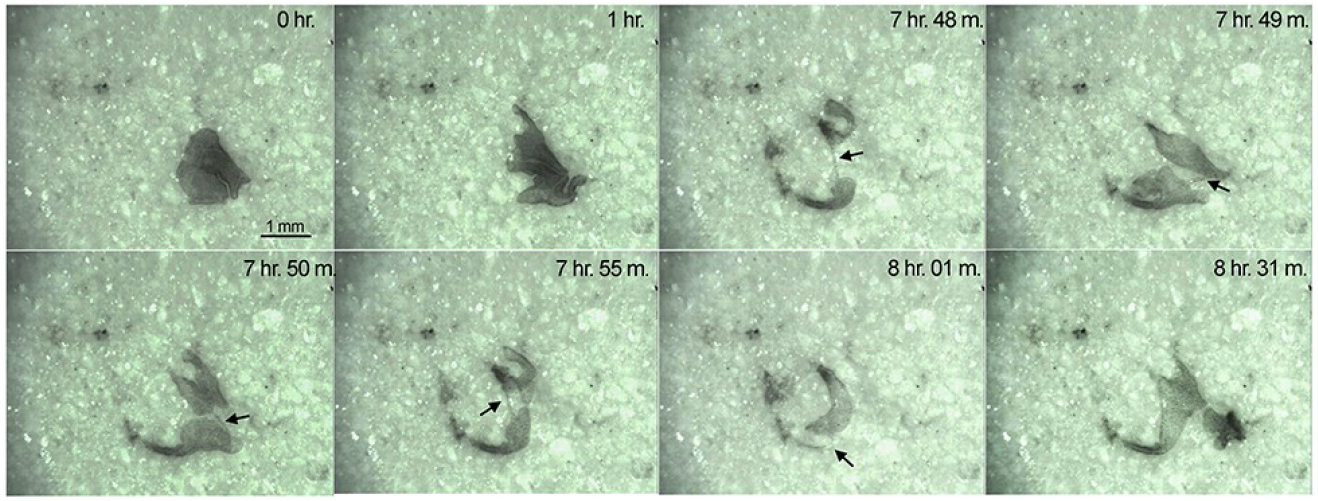
*Trichoplax* attached to the bottom of the dish (as a hard substrate) under a layer of kaoline; the time after sedimentation of the suspension is indicated on each picture. Frames “0 min” and “1hour” show the ability of *Trichoplax* to move in the sediment; the following frames after 7 hours show the fission of *Trichoplax* in two. The arrows mark the appearance of a thin cellular thread at the site of the fission and subsequent separation in two bodies (8 hours 01 min). See video 4.

In the second scenario, the *Trichoplax* dug itself out of the sediment and onto the surface. None of the animals originally placed in the cup (10 specimens) died, since after 18 hours they had all risen to the surface. Where they came to the surface, pits are visible in some cases; their walls seem to be reinforced with a polysaccharide film (Fig. 8A-C).

**Fig. 8.**
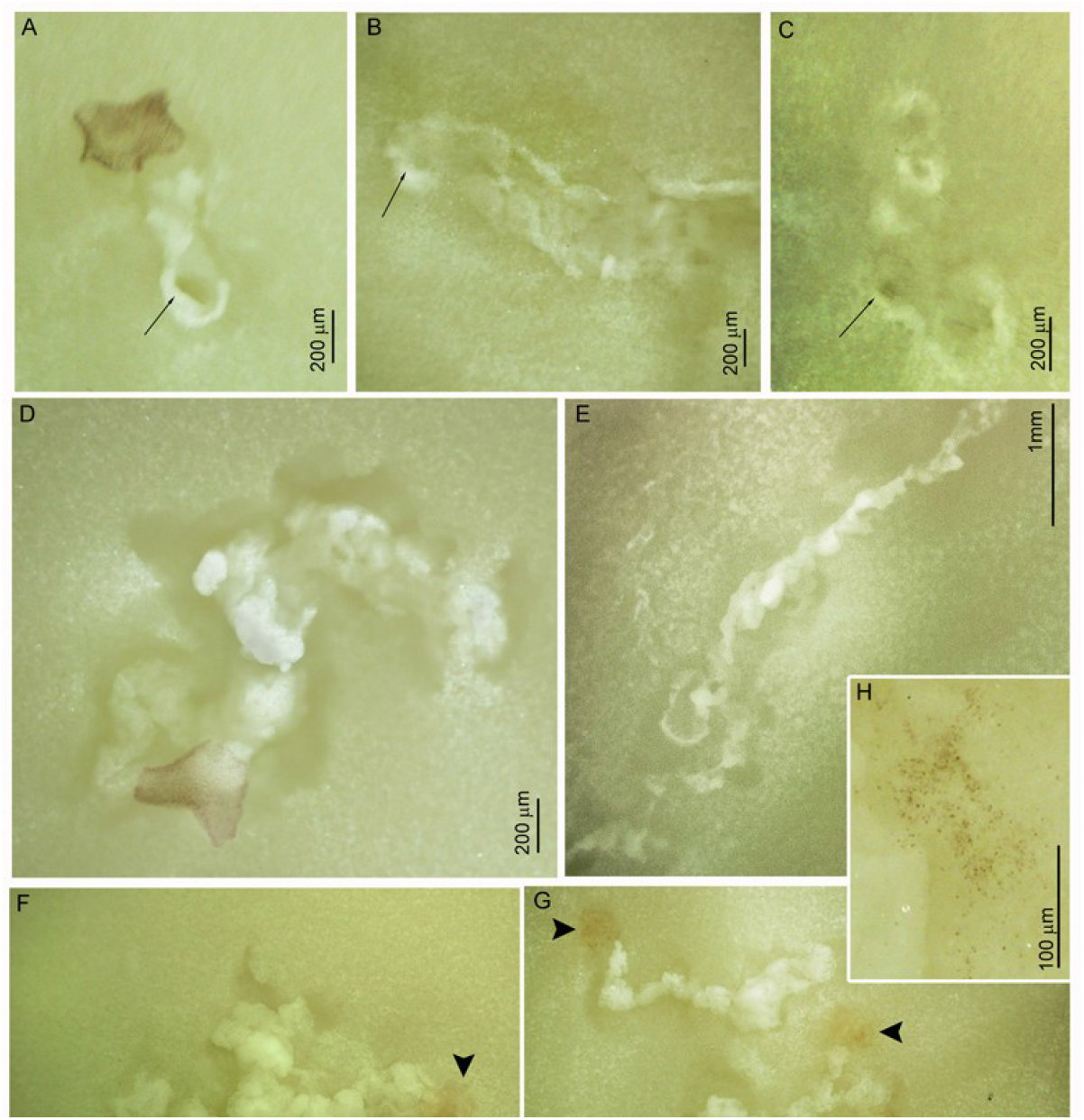
Traces of *T*.*adhaerens* movement on unconsolidated fine-grained sediment. A-C - funnel-shaped traces where *Trichoplax* creeped out onto the surface (long arrows). D, E, F - traces of the crawling of single individuals; D, F – wandering traces, E – straight trace, G – crawling traces of several individuals; the pigmented remnants of the dead producers are visible at the ends of each trace (short arrows). H - scattering of pigmented residues at the end of the trace.

### Traces of locomotion of *Trichoplax* on the loose sediment

After crawling out to the surface of the unconsolidated sediment or being simply placed on its surface, the animals began to move. Their traces and the way they moved bear little resemblance to the locomotion of *Trichoplax* on a solid surface.

Starting to move on the loose sediment, *Trichoplax* often consolidated a small area, and crawled several times around it. (See video 5). These launch grounds were larger in diameter than the size of its producers and the subsequent chain. Then the animal, moving a little to the side; compacted the next lump of sediment, and then moved away from this lump to the side a little more and compacted the next lump (videos 6, 7). Such locomotion resulted in the formation of chains of tightly packed lumps (Fig. 8D-G). These chains were three-dimensional, as the animals moved around the lumps from all sides: from above and below and from the sides. The lumps in the chains were smeared with a binding substance (most likely mucin secreted by the cells of the ventral cells), so they were relatively more stable than the surrounding sediment. The sizes of the lumps varied; in simple chains they never exceeded the area of its producer (Fig. 8E), and in the complex chains the lumps are combined in bigger clusters (Fig. 8F, G).

If several *Trichoplax*’s occurred together and they differed in size, then the larger individuals produced a trail chain, and the smaller ones simply used it for their own movement (See video 8). The animal could return to any point of its path and begin to form a new branch from the already existing chain. Two animals could connect their chains with a link. The resulting trails were as uneven curving chains with anastomoses and branches (Fig. 8F, G; video 8). Straight chains under the proposed experimental conditions were produced rarely (Fig. 8E).

After a day on the loose sediment, the animals inevitably died disintegrating into individual cells. Since in this experiment we used pigmented animals, we could see a spot of the pigmented dots on the sediment which are the individual cells (Fig. 8F, G, H). In no case did we observe a whole dead body, which would have stayed bound in one way or another with the sediment.

Given the possibility of fossilized “feeding” traces of Placozoans, we performed a corresponding experiment. To carry it out we grew bacterial mats on the kaoline sediment and on the fine-grained limestone sediment. On the kaoline mat, all specimens died within one hour. On the limestone mat, they moved actively for several hours however we did not observe any signs of the feeding behavior although the same mat on a hard surface, grown in parallel control, served as a good feeding ground for them.

## DISCUSSION

### The possibility of *Trichoplax* body fossils formation

Deposition of aluminum and silicon ions has been shown to facilitate the conservation and subsequent fossilization of soft-bodied organisms [19]. Conversely, adhesive complex molecules are responsible for the contact of cells with each other and with the substrate, and also most actively bind to Al [22]. Thus, Metazoans with a developed adhesive complex, when buried in aluminosilicate sediment, inevitably decay slowly after death and are therefore prone to form body fossils. In *Trichoplax*, this adhesive molecular complex is reduced, since it lacks claudins, occludins, and JAM proteins [7]. Yet, due to the presence of cadherins and integrins, the structure of cell contacts is similar to that of adhesive contacts (adherens junctions) in other Metazoa. In this regard, the preservation of the body remains of the *Trichoplax* should proceed via the same biochemical model as in other multicellular animals. Indeed, we have demonstrated that Al-cations can rapidly precipitate on the *Trichoplax* body remains promoting its preservation. Thus we can infer from this experiment that formation of placozoan body fossils is possible from the chemical view.

However, the postmortem transformations of *Trichoplax* have been shown to decrease this possibility to negligible values for two main reasons. The key obstacle to this is the almost instantaneous disintegration of a *Trichoplax* body into a cloud of individual cells. The corpses did not attach to solid (glass) or loose (clay) substrates, and dead cells rapidly floated away from the decaying/desintegrating body. Obviously, such a postmortem pattern precludes the formation of a recognizable multicellular body fossil. Even if the scattered cells were fossilized, a paleontologist could not claim to have found the remains of a multicellular organism. Related to our observation is the report about possible *Trichoplax*-like body and trace fossils from the Middle Triassic of Germany [27]. They are represented by small shapeless limonitic spots on the ends of some short slightly convex trails. They were buried under a rapidly deposited sediment layer as followed from their taphonomic description. The pattern contradicts with the results of our observations on the immediate after-death decay of Placozoa and on their resistance to the unfavorable burial factors. Thus, the assumed placozoan nature of these fossils needs to be proved by more facts and currently seems doubtful for us.

The high resistance to a lethal catastrophic burial lowers the probability of the formation of *Trichoplax*-like body fossils even more. During catastrophic events multicellular animals are buried under a thick sediment layer, and their bodies are quickly preserved as a result of the unfolding of specific physicochemical processes [18,19,21,28]. However, *Trichoplax* can exist under and inside fine-grained sediment as well as easily being able to get to the surface.

Conversely, the accumulation of hydrogen sulfide in the sediment is not too detrimental for *Trichoplax* as it can withstand moderate H_2_S contamination. Also, these animals can survive with a decreased proportion of oxygen respiration: they possibly have symbionts with other types of respiration or activate additional respiration mechanisms without cytochrome oxidase [29,30].

A slight acidification of sediment does not affect the viability of *Trichoplax* in any way. At borderline pH5.0, *Trichoplax* forms mobile spheres.

The high biochemical resistance of *Trichoplax* seems trivial. It is accounted for the effective homeostasis regulatory mechanisms, the presence of symbionts, as well as the legacy of unicellular ancestors that were richly equipped with various biochemical tools to support its-single-celled body in harsh chemical conditions. A non-trivial feature is the rapid post-mortem dissociation into cells. It occurs, apparently, due to the lack of a basement membrane underlying the cellular layer, as well as occluding (septate) cell-cell contacts in *Trichoplax* [31,32].

The basement membrane is present in all multicellular animals with the exception of Placozoa, Demospongia and ctenophores *Mnemiopsis leidyi*. Given the presence of the main components of the basement membrane (collagen IV and laminin) in all known Metazoa and the evolution of these proteins, the basement membrane has been proposed to be secondarily lost in those without it [32,33]. Nevertheless, *Trichoplax* can serve as a model for a metazoan in which the basement membrane has not yet been fully formed. According to modern ideas, such metazoan was a choanoblastaea, the hypothetical ancestor of multicellular animals [13,14,16,34].

Choanoblastaea has been assumed to be a sphere of collar cells connected by cell-cell contacts based on cadherins; integrins were involved in the binding of cells to an extracellular matrix, and tyrosine kinase mediated a cell-cell signaling. The early choanoblastaea did not yet have a basement membrane that united a layer of epithelial cells; only the late choanoblastaea developed it. It took a considerable time to evolve this membrane and occluding cell-cell contacts with the corresponding adhesive junction proteins, Neurexin IV, Claudin, and Contactin [16,31] which tightly zipped the cells together. Therefore, if the early choanoblastaea, like *Trichoplax*, could easily withstood harsh conditions and, without a basement membrane, disintegrated into single cells after death, it is impossible to trace this early stage in the fossil record.

We assume that the ancestors of metazoans could be biochemically quite robust, but their morphological stability was extremely ephemeral due to the absence of the basement membrane. This is probably one of the reasons for a gap between the appearance of multicellular organisms in the fossil record (about 600 million years ago) and the reconstructed molecular age of a common Metazoan ancestor (about 900-800 million years). The beginning of the fossil record of Metazoans may reflect the establishment of the basement membrane in their evolution; it tied the cells together even after the death of an animal, preserving its shape.

### The possibility of *Trichoplax* trace fossils

The tracks of *Trichoplax* movement on a solid surface are winding lines, as we can judge from the literature [35–37] and our observations. It should be emphasized that the characteristics of the tracks with and without chemotaxis, as well as their interspecific variability, are still poorly understood. However, the information available so far cannot be relied upon to identify Placozoan ichnofossils not because of the paucity of such data, but because tracks that may be preserved in the fossil record must be left on fine-grained sediment. Meanwhile, nothing has been found in the literature about how *Trichoplax* moves along loose sediment. It is probable that such a task has never been focused on; here it is presented for the first time to address the understanding of the early evolution of animals.

In experiments on the *Trichoplax* locomotion, we expected to see winding lines where clay particles were stuck together by mucus. However, the observed pattern turned out to be quite different. Once on soft sediment, *Trichoplax* formed a compacted launch mound, then lump after lump compacted a three-dimensional chain path as it advanced. The paths branched, since the animal could start a new chain from any place of the already formed chain, as well as connecting with the already existing paths of other individuals nearby. The nature of the movement – the chains of lumps, their branching and anastomoses turned out to be different to known placozoan tracks on a hard surface.

Such trails are knitted on all sides with mucus, an important component of which is mucin [10,36]. This acidic glycopolyprotein has a high affinity for positively charged cations and aluminosilicate particles, such that such a pathway may be preserved in the fossil record. It will have a three-dimensional configuration, so a paleontologist might easily mistake it for a burrowing trail, inferring the existence of burrowing organisms. The launch mounds at the beginning of the tracks differ in configuration from the rest of the chain, which also allows for a wide variety of misinterpretations about the morphology of a trail producer. Furthermore, branching and anastomosing are not characteristic of Placozoan tracks on a solid surface; this is important when deciding if problematic trace fossils are of a placozoan nature or not.

We suspect some resemblance between the *Trichoplax* traces and the Precambrian menisculate problematic ichnofossils. Menisculate ichnofossils usually look like short segments adjacent to each other in a chain-like manner. The segments are flattened, curved in the form of irregular arches or trapezoids, each arch is perpendicular to the bedding plane. The long or short chains of such meniscus-like segments form deep grooves in the bedding surface. Not uncommon are rounded flat spots or pits located within or at the beginning of these chains.

A variety of interpretations have been suggested for the menisculate ichnofossils: feeding traces, burrows, destroyed tubes, or some bodily structure that consisted of connected disks (review in [38]). For example, Rogov et al. [39], who focused on the low variability of width of each trail and their three-dimensional configuration in sediment, suggested that these were the earliest traces of burrowing multicellular animals. Accordingly, these authors associated the beginning of the substrate revolution with menisculate trails. The “burrowing hypothesis” contradicts the fact that the traces are present only on the bedding surface without signs of sediment mixing. Besides, the traces are often broken off and can be seen both as short chains and as separate segments. These observations allowed Ivantsov and colleagues to put forward a hypothesis about the “bodily” [40] or even skeletal [38] nature of menisculate *Nenoxites*, although the composition and structure of the presumable skeleton, as Ivantsov has admitted, has remained unknown. Since the traces of *Trichoplax* locomotion on loose sediment were not known, paleontologists had no reason to associate menisculate traces with placozoan-like locomotion.

*Trichoplax* moves according to the principle of soft adhesive locomotion [41,42], stretching out a part of its body, sticking it to the substrate, and pulling the other parts of its body over the mucus to the point of attachment. Unlike other multicellular adhesive movers, *Trichoplax* only uses cytoskeletons and cell-cell junctions in this locomotion but not muscles.

Such a mode of locomotion could have been found in early multicellular animals without muscles; here we mean choanoblastaea and its close descendants. If such non-muscular animals left their tracks on the sediment, then these tracks most likely looked quite different from the tracks known from muscular animals, for example, like the tracks that we observed when *Trichoplax* moved. At present, we have very little or no data on the footprints of animals without musculature.

Concerning the possible feeding traces in the fossil record, *Trichoplax* can hardly leave them on fine grained sediment. As we could see, *Trichoplax* obviously does not like to feed on a mat overgrown the loose sediment thus has little chance to left the feeding traces. Even if he fed on such a mat, he died very quickly. To explain this phenomenon, we need to recall how *Trichoplax* feeds. It creates a closed space between the ventral layer of cells and the surface of the substrate, where it releases digestive enzymes [9,43]. But loose – and especially aluminosilicate sediment – is an excellent adsorbent, and immediately absorbs the secreted enzymes. Thus *Trichoplax* wastes energy producing enzymes without getting anything in return (Fig. 8F, G). We also can suppose that something was released through the interaction between bacteria and sediment unsuitable for *Trichoplax* survival. According to field observations, *Trichoplax* seemingly avoid such substrates: it live and can be caught above a rocky bottom, but they are extremely rarely caught above a soft substrate [44,45].

The Placozoan feeding mode has been proposed to be similar to the early animals such *Dickinsonia* [46,47]. The provisional feeding traces of *Dickinsonia* were formed on unconsolidated (possibly clay) sediment overgrown with a microbial mat. These traced have the shape of *Dickinsonia* ventral (?) surface and were numerous on the bedding plane. We have shown that such fossils would unlikely be formed via the Placozoan mode of feeding. Thus we supposed that *Dickinsonia* differed from the known extant placozoans in their feeding behavior. However, we still cannot imagine what external digestion looks like in *Dickinsonia* and other animals of this kind, that is, with a basal lamina, since they did not lose their shape after death, but without a gut [47].

## CONCLUSIONS

1. *T. adhaerens* has shown considerable biochemical stability and a possibility to survive in sediment. After death, their bodies immediately disintegrate to a cloud of individual cells. Thus the *Trichoplax*-like body remains have little chances for preservation and fossilization.
2. We hypothesized that the immediate after-death dissociation of such a body into cells is accounted for the absence of a basal membrane underlying epithelial layer, while the presence of the basal membrane provides preservation the shape of an animal after its death. Consequently, the evolutionary development of the basal membrane in Metazoa seems to become a starting point for their fossil record.
3. Traces of *Trichoplax* locomotion on loose sediment appeared to have unusual configuration previously unknown: these are three-dimensional chains of substrate lumps, densely packed and adjacent to each other. Besides, they have branches and anastomoses. These tracks strikingly differ from those on solid substrates. Given that traces could be preserved and fossilized namely in unconsolidated deposits, fossil traces of the placozoan-like locomotion should remind such chains. We suggested that some early menisculate ichnofossils could be produced by a placozoan type of locomotion.
4. The feeding traces of *Trichoplax* on an unconsolidated substrate overgrown by a bacterial mat do not form (or were not observed) because of their inability to feed on loose sediment. Thus the feeding traces of the Placozoan-like feeding behavior would unlikely preserved in fossil record.

## ACKNOWLEDGEMENTS

The work of YVL and ADF was carried out using the equipment of the Center for Collective Use of the Institute of Developmental Biology. N.K. Koltsov of the Russian Academy of Sciences (IBR RAS) with the support of the State Assignment of the IBR RAS (No. GZ 0088-2021-0008).

## CAPTIONS FOR THE VIDEOS

Video 1. Post-mortem changes of *Trichoplax* at the lethal concentration of sodium azid (0,4%, see Fig. 4); the body contracted, then single cells detached from the outer edge following by an abrupt bursting into a cloud of dead (pink) cells.

Video 2. Post-mortem changes of *Trichoplax* under UV radiation; it moved very fast for the first 10 min, then contracted and in one minute, burst into individual cells which started floating away and continued this for the next 40 minutes. Staining by DAPI with the nuclei in bright blue; note light-blue nuclei-free remnants in the expanding cloud of dead cells.

Video 3. A Trichoplax sphere had formed at pH 6.0 and then exposed in azide (0.4%): it immediately disintegrated into single cells.

Video 4. *Trichoplax* can survive, actively moves, and divides under the layer of kaoline sediment (3mm thick); this fragment starts after 7 hours after the beginning of the experiment and includes a moment of the division.

Video 5. The starting sites of two trails: Two *Trichoplax* specimens move around the sites to consolidate their starting grounds.

Video 6. The trace of the *Trichoplax* locomotion on kaoline; example 1.

Video 7. The trace of the *Trichoplax* locomotion on kaoline; example 2

Video 8. Several *Trichoplax* move: the bigger ones make the trail while the smaller ones use it for their locomotion. The *Trichoplax* specimens can join their trials producing anastomoses or return to any site of the trail and make a branch (the right trail).

